# Endurance exercise with reduced muscle glycogen content influences substrate utilization and attenuates acute mTORC1- and autophagic signaling in human type I and type II muscle fibers

**DOI:** 10.1101/2024.10.14.618241

**Authors:** O Horwath, L Cornet, H Strömlind, M Moberg, S Edman, K Söderlund, A Checa, J L Ruas, E Blomstrand

**Author notes:** Corresponding author Eva Blomstrand Department of Physiology, Nutrition and Biomechanics, The Swedish School of Sport and Health Sciences, Stockholm, Sweden.

## Abstract

**Background:** Exercising with low muscle glycogen content can improve training adaptation, but the mechanisms underlying the muscular adaptation are still largely unknown. In this study, we measured substrate utilization and cell signaling in different muscle fiber types during exercise and investigated a possible link between these variables.

**Methods:** Five subjects performed a single leg cycling exercise in the evening (day 1) with the purpose of reducing glycogen stores. The following morning (day 2), they performed two-legged cycling at ∼70% of VO2peak for 1h. Muscle biopsies were taken from both legs pre- and post-exercise for enzymatic analyses of glycogen, metabolite concentrations using LC-MS/MS-based quantification, and protein signaling using Western blot in pools of type I or type II fibers.

**Results:** Glycogen content was 60-65% lower for both fiber types (P<0.01) in the leg that exercised on day 1 (low leg) compared to the other leg with normal level of glycogen (normal leg) before the cycling exercise on day 2. Glycogen utilization during exercise was significantly less in both fiber types in the low compared to the normal leg (P<0.05). In the low leg, there was a 14- and 6-fold increase in long-chain fatty acids conjugated to carnitine in type I and type II fibers, respectively, post-exercise. This increase was 3-4 times larger than in the normal leg (P<0.05). Post-exercise, mTOR^Ser2448^ phosphorylation was increased in both fiber types in the normal leg (P<0.05) but remained unchanged in both fiber types in the low leg together with an increase in eEF2^Thr56^ phosphorylation in type I fibers (P<0.01). Exercise induced a reduction in the autophagy marker LC3B-II in both fiber types and legs, but the post-exercise level was higher in both fiber types in the low leg (P<0.05). Accordingly, the LC3B-II/I ratio decreased only in the normal leg (75% for type I and 87% for type II, P<0.01).

**Conclusions:** Starting an endurance exercise session with low glycogen availability leads to profound changes in substrate utilization in both type I and type II fibers. This may reduce the mTORC1 signaling response, primarily in type I muscle fibers, and attenuate the normally observed reduction in autophagy.

## Background

Endurance exercise performed with low levels of muscle glycogen influences not only performance, but also substrate utilization and protein metabolism, and has been reported to improve the muscular adaptation to training [1–4]. The rate of glycogen utilization is lower, glucose uptake is higher, and there is a significant net protein degradation during exercise even when delivery of blood-borne substrates and hormones are the same to both legs [5–7].

The rate of protein synthesis is regulated by the mechanistic target of rapamycin complex 1 (mTORC1) and subsequent activation of the downstream effector proteins p70 ribosomal protein S6 kinase 1 (S6K1), the eukaryotic initiation factor 4E-binding protein (4E-BP1) and the eukaryotic elongation factor 2 (eEF2) [8, 9]. There is some evidence that the rate of protein synthesis decreases during aerobic exercise [10], which is reflected in elevated eEF2 and reduced 4E-BP1 phosphorylation [11], and then increases again during recovery to levels higher than before exercise [12–14].

Furthermore, the major systems involved in protein degradation are the ubiquitin-proteasome system and the autophagy-lysosome pathway [15]. Molecular markers for the former pathway, atrogin-1 and muscle ring-fiber protein-1 (MuRF-1), are upregulated at the mRNA level following a session of endurance exercise [16]. The autophagy pathways appear to be intensity-dependent, activated by high-intensity exercise via the activation of AMP-activated protein kinase (AMPK) and its downstream protein unc-51 like autophagy-activating kinase-1 (ULK1), whereas endurance exercise of moderate intensity decreases the autophagosome content [17, 18].

Little is known about the impact of exercising with low glycogen levels on signaling pathways regulating muscular adaptation [19–21]. In addition, these studies are commonly conducted on mixed muscle, whereas the adaptive response is likely to be influenced by the type of fibers that are activated [22]. A difference between the major muscle fiber types with regard to the expression of genes regulating muscle metabolism and protein signaling, as well as a differential response to both endurance and resistance exercise, have previously been reported [23–27]. However, no studies have investigated a possible fiber type-specific response to aerobic exercise with limited availability of muscle glycogen. During this exercise, the recruitment of fibers may be changed compared to a normal nutritional condition. Measurements of substrate utilization and cell signaling in the various fiber types may therefore provide novel insights into muscle physiology that are overlooked using conventional analyses on biopsies from whole muscle.

The aim of the study was therefore to investigate the effect of endurance exercise with reduced initial glycogen availability on muscle metabolism and cell signaling in type I and type II fibers. With this purpose, subjects performed one-legged cycling exercise in the evening to reduce the muscle glycogen content. The following morning, they performed cycling exercise with both legs. Muscle biopsies were obtained before and after the morning exercise for analyzes of substrates and metabolites as well as cell signaling in pools of type I and type II fibers. We hypothesized that in a muscle that performs exercise with reduced muscle glycogen availability, markers for anabolic and catabolic pathways would be changed in such a way as to promote muscle protein degradation in the muscle that begins the exercise with reduced muscle glycogen, mainly in type II fibers because of their greater reliance on glycogen as a substrate.

## Material and methods

### Subjects

Five healthy subjects (4 males and 1 female) participated in the study. They were all moderately trained, performing endurance and/or resistance exercise 3-4 times per week. Their mean (± standard error (SE)) age was 25 (± 1) years, height 183 (± 3) cm, body mass 74 (± 5) kg and maximal oxygen uptake (VO2peak) 3.89 (± 0.36) l min^-1^. All participants were fully informed about the experimental procedure and associated risks before giving their written consent. The study was approved by the Swedish Ethical Review Authority (2018/2186-31) and performed in accordance with the principles outlined in the Declaration of Helsinki.

### Experimental design

#### Preliminary tests

The preliminary exercise tests were performed on a mechanically braked cycle ergometer (Monark 828E, Vansbro, Sweden). One week before the experiment, the oxygen uptake of the subjects was determined at three submaximal work rates, along with their peak oxygen uptake (VO2peak) using an on-line system (Oxycon Pro, Jaeger, Hoechberg, Germany). The subjects exercised at a pedaling rate of 70 rpm. A work rate corresponding to approximately 70% of VO2peak was calculated from these measurements.

#### Exercise for glycogen reduction (day 1)

A schematic overview of the experimental workflow is provided in Figure 1. During the two days preceding the experiment, subjects were instructed to refrain from intensive exercise and keep a record of their dietary intake. The subjects came to the laboratory between 5 and 6 PM on the evening before the experiment. They performed one-legged cycling on a mechanically braked ergometer (Monark 828E) equipped with a custom-made pedal that held a 5 kg counterweight to assist with the upward phase of the pedaling action [28, 29], with the other leg resting on a chair. The exercising leg was randomly selected. The exercise protocol has been described in detail previously [30], and an overview of this protocol is provided in Figure 2. Briefly, the subjects performed one-legged cycling for 45 min at 80 rpm at a work rate of 111 ± 12 W (heart rate 144 ± 4 beats per minute (bpm)). After a 5 min rest, they then performed interval exercise, consisting of 5 x 2 min one-legged cycling at a work rate of 149 ± 16 W. This was followed by an interval exercise of 5 x 3 min of maximal arm cranking using both arms (Monark 891E Wingate) at a work rate of 94 ± 16 W. The protocol was designed to lower the glycogen content in both type I and type II fibers, and by adding arm exercise the rate of glycogen resynthesis in the exercised leg during the rest period until the following morning [30]. After this exercise session, the subjects remained fasted until the experiment the next morning.

**Figure 1.**
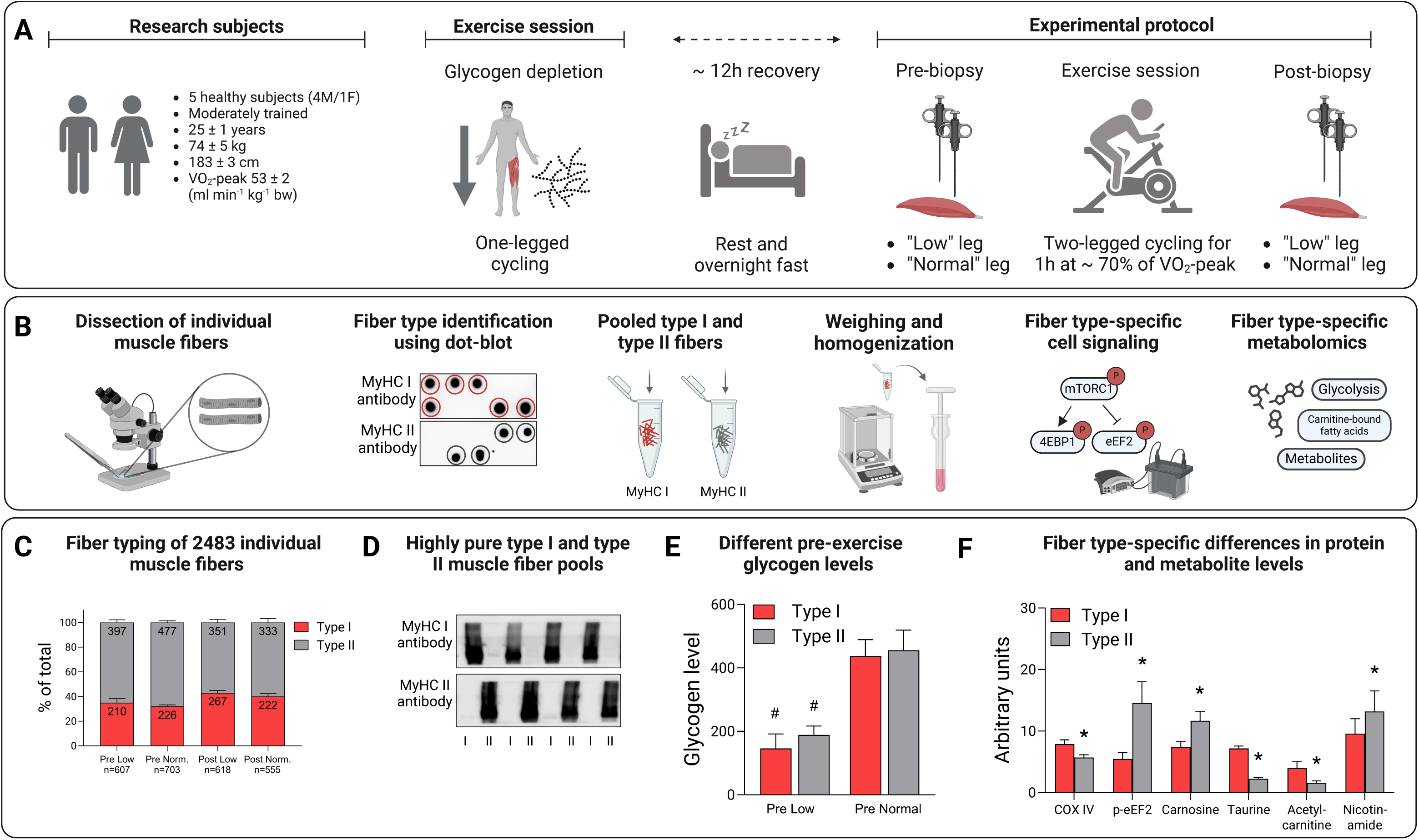
Workflow of the one- and two-legged exercise and muscle fiber isolation, identification, and analysis. A. Subject characteristics and schematic description of the exercise protocol, B. Individual fibers were separated from freeze-dried muscle biopsies, classified by dot-blotting and pooled into groups of type I and type II fibers. The fiber pools were weighed and homogenized in western blot buffer (∼ 1µg ml µl^-1^) and analyzed with regard to proteins in the mTORC1 pathway as well as substrate and metabolites using metabolomics, C. Number and proportion of type I and type II fibers dissected out from biopsies obtained from the low and normal leg pre- and post-exercise, D. The purity of type I and type II fiber pools confirmed by analyzing the homogenates with antibodies against MyHC I or MyHC II, E. The one-legged exercise resulted in a 60-65% reduction in muscle glycogen the following morning, ^#^P<0.001 for low versus normal leg, and F. Fiber-type specific differences in protein and metabolite levels in biopsies from the normal leg pre-exercise, *P<0.05 for type II versus type I fibers.

**Figure 2.**
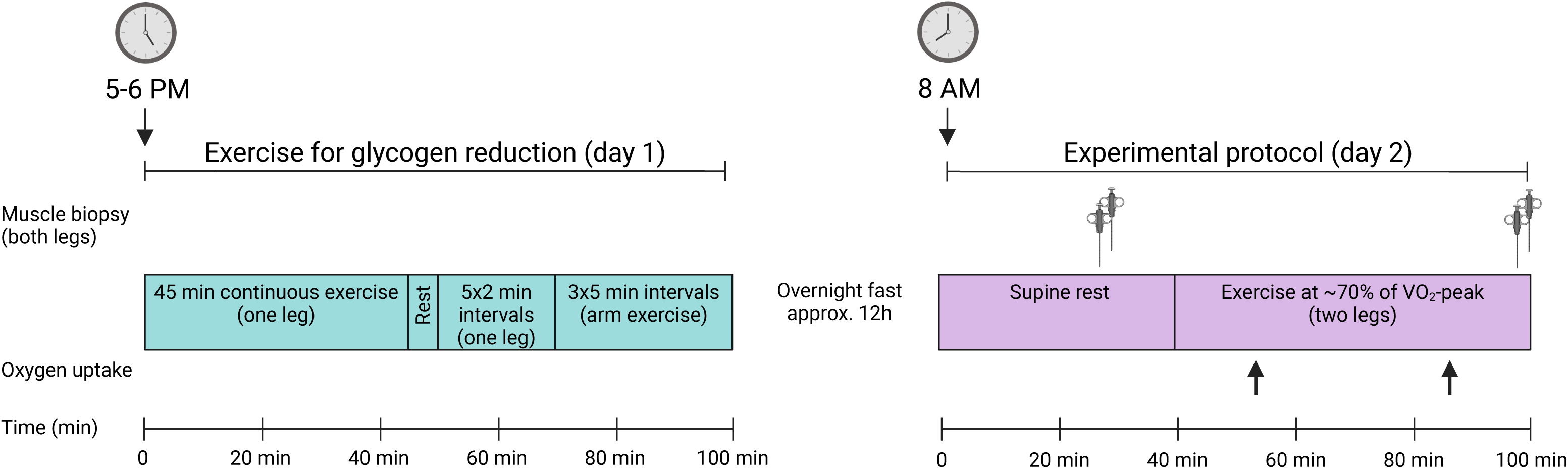
A schematic overview of the one-legged exercise session on the evening of day 1 performed to reduce glycogen levels, and the two-legged exercise session performed in the morning of day 2.

#### Experimental protocol (day 2)

The subjects reported to the laboratory in the morning at 8 AM after fasting overnight. After 30 min supine rest, muscle biopsies were taken from the vastus lateralis (∼ 10 cm from patella) of both legs under local anaesthesia (2% Carbocain, Astra Zeneca, Södertälje, Sweden) using a Weil-Blakesley conchotome (AB Wisex, Mölndal, Sweden) as described by Henriksson (1979) [31].

The subjects then exercised for 60 min at 70 rpm on a cycle ergometer (Monark 828E) at a work rate corresponding to ∼ 70% of VO2peak, see Figure 2. Pulmonary oxygen uptake was measured after 10-15 and 40- 45 min of exercise using an on-line system (Oxycon Pro, Jaeger) and heart rate was monitored continuously by a portable device (Polar Electro Oy, Kempele, Finland). The relative pedal force of each leg was recorded during exercise using a pedal-based power meter system (Vector 2, Garmin, Kansas USA), and the data were processed with publicly available cycling performance software (GoldenCheetah, version 3.3). Immediately after exercise the subjects lay down and a second muscle biopsy was taken from both legs using the same procedure as pre-exercise. The second biopsy was taken 2-3 cm proximal to the first one and both biopsies were taken at approximately the same position in both legs. The muscle samples were freed from blood and quickly frozen in liquid nitrogen and stored at - 80 °C.

#### Single fiber dissection and type identification

The muscle samples were freeze-dried overnight and subsequently dissected free of blood and connective tissue. Thereafter, single fibers were dissected out using needles and a fine forceps under a stereo microscope (Carl Zeiss MicroImaging, Jena, Germany).

In order to determine muscle fiber type, one fragment of the isolated single fibers was cut off and dissolved in 5 µl western blot (WB) buffer (2 mM HEPES (pH 7.4), 1 mM EDTA, 5 mM EGTA, 10 mM MgCl2, 1% Triton X-100, 1 mM Na3VO4, 2 mM dithiothreitol, 1% phosphatase inhibitor cocktail (Sigma P- 2850) and 1% (vol/vol) Halt Protease Inhibitor Cocktail (Thermo Scientific, Rockford, IL)) and 5 µl 2X Laemmli sample buffer (Bio Rad Laboratories, Richmond, CA) and heated at 95°C for 5 min. The fibers were then identified as either type I or type II using a modified version of the dot blotting procedure described by Christiansen et al. (2019) [32]. Briefly, polyvinylidene fluoride (PVDF) membranes were activated in 95% ethanol for 15–60 s and then equilibrated for 2 min in transfer buffer (25 mM Tris, 192 mM glycine, pH 8.3 and 20% methanol). 1 µl of each sample was applied to a specific part of two membranes. After complete absorption of samples, the membrane was left to dry for 2–5 min before being reactivated in 95% ethanol for 15–60 s and equilibrated in transfer buffer for 2 min. After washing in Tris-buffered saline-Tween (TBST), the membranes were blocked in 5% non-fat milk in TBST (blocking buffer) for 5 min at room temperature.

Following blocking, the membranes were rinsed with TBST and then incubated with antibodies against MyHC I (Abcam #ab11083, diluted 1:10,000) or MyHC II (Abcam #ab91506, diluted 1:10,000) at room temperature for 2 h. Membranes were washed and incubated with secondary antibodies, anti-mouse (Cell Signaling Technology #7076S; 1:10,000) or anti-rabbit (Cell Signaling Technology #7074; 1:10,000) for 1 h at room temperature followed by washing in TBST. Proteins were visualized by applying Super Signal West Femto Chemiluminescent Substrate (Thermo Scientific) to the membranes, followed by detection on a Molecular Imager ChemiDoc™ MP system. A representative picture of the dot-blot image is provided in Figure 1B.

### Pooling of single fibers

Based on the dot blot, fibers were classified and pooled into groups of type I and type II fibers. The pools of fibers were then weighed on a Cubis^®^ high-capacity micro balance (Sartorius Lab Instruments, Göttingen, Germany). The average weight of the fiber pools was 144.1 µg (range 33.3–359.8 µg). The fiber pools were homogenized in ice-cooled WB buffer using a ground glass homogenizer to a concentration of ∼ 1 µg/µl, except in the case of the low weight pools where the concentration was less, and stored at -80 °C.

### Analysis of fiber pools

#### Determination of glycogen content

The glycogen concentration in the homogenate was measured according to the method described by Leighton et al. (1989) [33]. 10 µl homogenate was digested in 1 M KOH at 70 °C for 15 min. After cooling, pH was adjusted to 4.8 with glacial acetic acid followed by addition of 10 µl acetate buffer (pH 4.8) containing amyloglucosidase and incubated at 40 °C for 2 h. Glucose concentration was then analyzed photometrically in a plate reader (Tecan Infinite F200, Männedorf, Switzerland). All samples were analyzed in triplicate.

### Metabolomic analysis

On the day of analyses, 10 µl of sample homogenates and blanks (homogenizing buffer) were reconstituted in 400 μl of LC-MS methanol, vortexed for 10 s and sonicated for 15 min on ultrasound bath on ice. Samples were then centrifuged at 10,000 *g* for 15 min. Finally, 80 μl were transferred to an LC-MS vial equipped with a 150 μl insert. Samples were injected in randomized order of fiber type within the same individual.

### LC-MS/MS analysis

Samples were analyzed on an ACQUITY UPLC System coupled to a Waters Xevo® TQ-S triple quadrupole system (both from Waters Corporation (Milford, MA)), equipped with an electrospray ion source. For all metabolites, at least one specific selected reaction monitoring (SRM) transition was analyzed. Two independent injections were performed in positive and negative ionization mode. In positive mode, metabolites were separated on an Acquity Premier BEH Amide Vanguard FIT column (100 x 2.1 mm, 1.7 µm). Aqueous mobile phase (MPA) consisted of 20 mM ammonium formate + 0.1% formic acid in double-deionized water. Organic mobile phase (MPB) consisted of 0.1% formic acid in acetonitrile. The following chromatographic gradient was used: 0 min, 95% B; time range 0 → 1.5 min, 95% B (constant); time range 1.5 → 14 min, 95 → 55% B (linear decrease); time range 14 to 14.2 min, 55 → 45% B (linear decrease); time range 14.2 → 16.0 min, 45% B (isocratic range); time range 16.0 → 16.2 min, 45 → 95% B (linear increase). The column was then equilibrated at 95% B for 7 additional minutes. The flowrate was 400 µl/min and the column temperature was held at 30°C. The volume of injection was set at 2.5 µl.

In negative mode, metabolites were separated on an Acquity Premier BEH Z-HILIC Vanguard FIT column (100 x 2.1 mm, 1.7 µm). Aqueous mobile phase (MPA) consisted of 20 mM ammonium acetate in double-deionized water adjusted to pH 9.1 with acetic acid. Organic mobile phase (MPB) consisted of acetonitrile. The following chromatographic gradient was used: 0 min, 90% B; time range 0 → 6 min, 90 → 65% B (linear decrease); time range 6.0 → 7.0 min, 65 → 55 %B (linear decrease); time range 7 to 7.5 min, 55 → 45% B (linear decrease); time range 7.5 → 9.5 min, 45% B (isocratic range); time range 9.5 → 10 min, 45 → 90% B (linear increase). The column was then equilibrated at 90% B for 4 additional minutes. The flowrate was 500 µl/min and the column temperature was held at 30°C. The volume of injection was set at 2.5 µl.

### Western blotting

Aliquots of the homogenates were diluted with 4X Laemmli sample buffer (Bio-Rad Laboratories) to a concentration of 0.25 µg muscle/µl and heated at 95°C for 5 min. Proteins were separated by SDS-PAGE, 15 µl from each sample was loaded onto 26-well Criterion TGX gradient gels (4–20% acrylamide, Bio Rad Laboratories) and electrophoresis run as previously described [34, 35]. Proteins were transferred to PVDF membranes (Bio Rad Laboratories) and stained with MemCode Reversible Protein Stain Kit (Thermo Scientific). All samples from each subject were loaded onto the same gel and all gels were run simultaneously. After destaining, the membranes were blocked in Tris-buffered saline (TBS; 20 mM Tris base, 137 mM NaCl, pH 7.6) containing 5% nonfat dry milk for 1 h at room temperature and incubated overnight with primary antibody. After incubation with primary antibody, membranes were washed and incubated with a secondary anti-rabbit or anti-mouse antibody for 1 h at room temperature followed by washing in TBST. Proteins were visualized by applying Super Signal West Femto Chemiluminescent Substrate (Thermo Scientific) to the membranes, followed by detection on a Molecular Imager ChemiDoc™ MP system and quantification of the resulting bands with Image Lab™ Software (Bio-Rad Laboratories).

The purity of the type I and type II fiber pools were confirmed by analyzing the homogenates as described above and incubated with antibodies against MyHC I (Abcam #ab11083, diluted 1:10,000) or MyHC II (Abcam #ab91506, diluted 1:10,000). A representative immunoblot is presented in Figure 1D.

### Antibodies

For immunoblotting, primary antibodies against mTOR (Ser^2448^, #2971; total, #2983), 4E-BP1 (Thr^37/46^, #2855; total, #9644), eEF2 (Thr^56^, #2331; total, #2332), S6K1 (Thr^389^, #9234; total #2708, ACC (Ser^79^, #3661; total, #3676), AMPK (Thr^172^, #4188; total, #2532), ULK1 (Ser^555^, #5869; total, #8054), LC3B (#2775), and COX IV (#4860) were purchased from Cell Signaling Technology (Beverly, MA, USA). Primary antibody for total MuRF-1 (#sc-398608) and UBR5 (#sc-515494) were purchased from Santa Cruz Biotechnology (Heidelberg, Germany).

All primary antibodies were diluted 1:1,000 except for total eEF2 and total 4E-BP1, which were diluted 1:2,000, and total UBR5 and MuRF-1 which were diluted 1:500. Secondary anti-rabbit (#7074; 1:10,000) and secondary anti-mouse (#7076; 1:10,000) were purchased from Cell Signaling Technology.

### Statistics

Conventional methods were employed to calculate means and SE of the mean. A three-way repeated measures ANOVA was employed to compare substrate, metabolite, and protein levels in type I and type II fibers in the low- and normal-glycogen leg pre-exercise as well as changes during exercise in both fiber types in the low- and normal-glycogen leg (leg, fiber type, time). When a significant main effect and/or interaction was observed in the ANOVA, Fisheŕs LSD post hoc test was employed to identify where these differences occurred. For variables with missing values, linear mixed-effects models were created with protein levels and interactions as fixed effects, while variation between individuals was treated as random effects. The emmeans package was used to estimate marginal means between leg, time and fiber type.

Pearsońs correlation coefficient (r) was calculated to evaluate a possible relationship between parameters analyzed in the muscle biopsies.

Statistical analyses were performed in Statistica version 13 (StatSoft Inc., Tulsa, USA) and in R version 4.3.0 with a P-value < 0.05 being considered statistically significant. Figures were created using GraphPad Prism version 9.1.2 for Windows (GraphPad Software, San Diego), with the exception of Figures 1 and 2 that were created with BioRender.com.

## Results

### Effects of the glycogen reduction exercise on pre-exercise variables (day 2)

#### Substrates and metabolites

Glycogen content in type I and type II fibers in the leg with reduced muscle glycogen (low leg) was 35% and 40%, respectively, of the content in the leg with normal glycogen level (normal leg) (P<0.001 for both fiber types, Figure 3A). The levels of the glycolytic metabolites glucose-6-phosphate (G-6-P) and fructose-6- phosphate (F-6-P) were three times higher in type II compared to type I fibers (P<0.05) in the low leg (Figure 3B, C). The levels of fatty acids conjugated to carnitine did not differ between conditions or fiber types.

**Figure 3.**
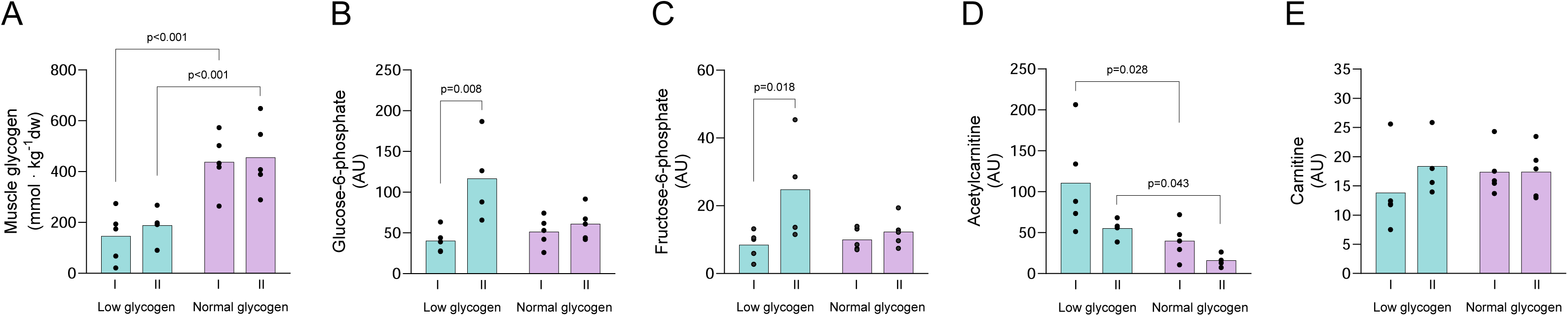
Levels of substrates and metabolites in type I and type II fibers in the low and normal glycogen leg pre-exercise on day 2. (A) muscle glycogen, (B) glucose-6-phosphate, (C) fructose-6-phosphate, (D) acetylcarnitine, and (E) carnitine. Individual values from 5 subjects as well as mean values are presented, except for type II fibers in the low leg pre-exercise and type I fibers in the normal leg post exercise where n=4 in figures B to E. A 3-way ANOVA was employed (A) or linear mixed effects models for variables with missing values (B-E) to compare levels in the low and normal leg as well as between fiber types.

Acetylcarnitine was considerably higher in both type I (2.8-fold) and type II fibers (3.4-fold) in the low leg (P<0.05 for both fiber types), whereas carnitine levels did not differ significantly between the low and normal leg (Figure 3D, E).

#### Cell signaling

The level of mTOR^Ser2448^ phosphorylation was significantly elevated pre-exercise in both fiber types (150% higher in type I and 75% in type II) in the low compared to the normal leg (P<0.01 for type I and P<0.05 for type II fibers) (Figure 4A). The phosphorylated levels of 4E-BP1^Thr37/46^, eEF2^Thr56^, AMPK^Thr172^, acetyl-CoA carboxylase (ACC^Ser79^) and ULK1^Ser555^ were similar in the low and normal leg. S6K1^Thr389^ phosphorylation was not measurable due to weak antibody signal.

**Figure 4.**
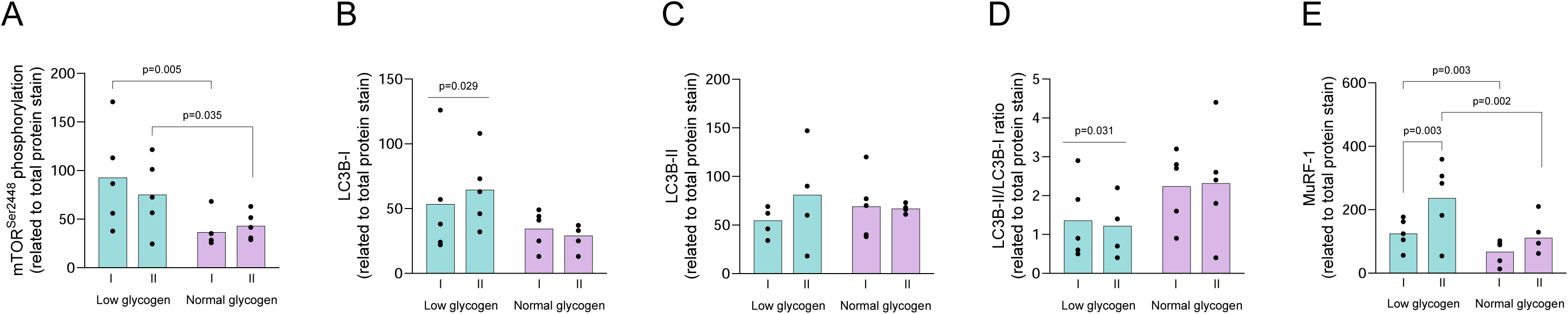
Levels of phosphorylated (p) or total protein of enzymes involved in synthesis and breakdown in type I and type II fibers in the low and normal glycogen leg pre-exercise on day 2. (A) p-mTOR, (B) LC3B-1, (C) LC3B-II, (D) LC3B-II/I ratio and (E) MuRF-1. Individual values from 5 subjects as well as mean values are presented. When the ANOVA revealed a significant interaction between leg and time or between leg, fiber type and time, Fisheŕs LSD post hoc test was employed to identify where the differences occurred.

The content of the autophagy protein LC3B-I was significantly higher in the low compared to the normal leg (P<0.05), 55% higher in type I and 120% in type II fibers, whereas the content of LC3B-II was similar in both legs (Figure 4B, C). Accordingly, the LC3B-II/LC3B-I ratio in the low leg was reduced by 61% and 53% in type I and type II fibers, respectively (P<0.05 for low vs. normal leg), with no difference between the fiber types (Figure 4D). The expression of the ubiquitin ligase MuRF-1 was significantly upregulated in both fiber types, although to larger extent in type II fibers (56% in type I and 123% in type II) in the low leg (P<0.05 for leg and fiber type, Figure 4E).

### Effects of two-legged exercise with reduced and normal muscle glycogen content (day 2)

#### Physiological parameters

All subjects completed the 60 min cycling exercise at an average work rate of 197 ± 21 W. Oxygen uptake during exercise averaged 2.83 ± 0.29 l min^-1^, corresponding to 73 ± 1% of VO2peak, respiratory exchange ratio (RER) averaged 0.87 ± 0.01, and heart rate 159 ± 4 bpm. The relative force produced by the low and normal legs when pedaling averaged 51 ± 1% for the low and 49 ± 1 % for the normal leg, with no significant difference between the two.

#### Muscle metabolism

The cycling exercise reduced the muscle glycogen level in both fiber types in both legs. However, the decrease was larger in the normal than in the low leg for both fiber types (Figure 5A). Glycogen content in type I fibers decreased from 146 ± 45 to 77 ± 42 mmol/kg dry weight (dw) in the low (P<0.05) and from 438 ± 51 to 92 ± 37 mmol/kg dw in the normal leg (P<0.001). In type II fibers, glycogen content decreased from 189 ± 28 to 90 ± 48 mmol/kg dw in the low (P<0.01) and 456 ± 63 to 225 ± 86 mmol/kg dw in the normal leg (P<0.001). The rate of glycogen utilization did not differ between the two fiber types in the low leg, whereas in the normal leg, the rate was 50% higher in type I than in type II fibers (P<0.01 for type I vs. type II) (Figure 5A). The decrease in glycogen during exercise was positively correlated to the initial content of glycogen in the type I fibers, whereas no such correlation was found in type II fibers (Figure 5B, C).

**Figure 5.**
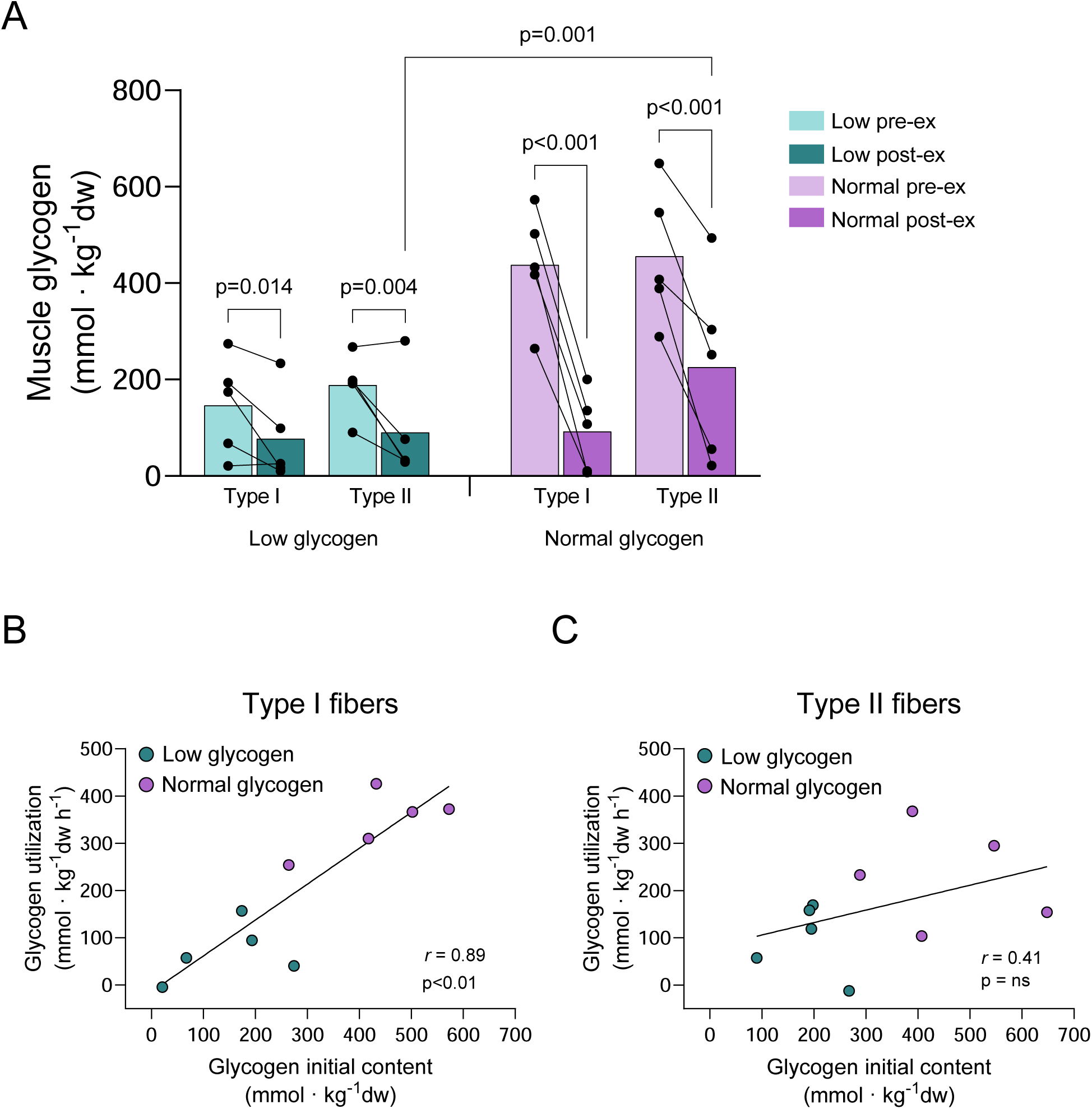
(A) muscle glycogen in type I and type II fibers in the low and normal glycogen leg pre and post 60 min of two-legged cycling exercise, (B) correlation between initial content of glycogen and utilization of glycogen in type I fibers and (C) corresponding correlation in type II fibers. Individual values from 5 subjects as well as mean values are presented. When the ANOVA revealed a significant interaction between leg, fiber type and time, Fisheŕs LSD post hoc test was employed to identify where the differences occurred.

Exercise led to pronounced increases in carnitine-conjugated long-chain fatty acids, namely, a 14-fold increase in type I fibers (P<0.001), and a 6-fold increase in type II fibers in the low leg (P<0.001, Figure 6A shows palmitoyl-carnitine). In the normal leg, there was a 7-fold increase in type I fibers (P<0.05), but no change in the type II fibers after exercise (Figure 6A). The increase in palmitoyl-carnitine was related to the initial muscle glycogen level in an inverse curve linear way (Figure 6B). A similar pattern of exercise-induced change was found for medium-chain fatty acids conjugated to carnitine (Figure 6C shows decanoyl-carnitine), whereas short-chain fatty acids conjugated to carnitine increased similarly in type I and type II fibers in both legs, although the increase was larger in the low leg (Figure 6D shows propionyl-carnitine). The remaining long, medium, and short-chain fatty acids conjugated to carnitine that were analyzed are presented in Supplementary figure 1.

**Figure 6.**
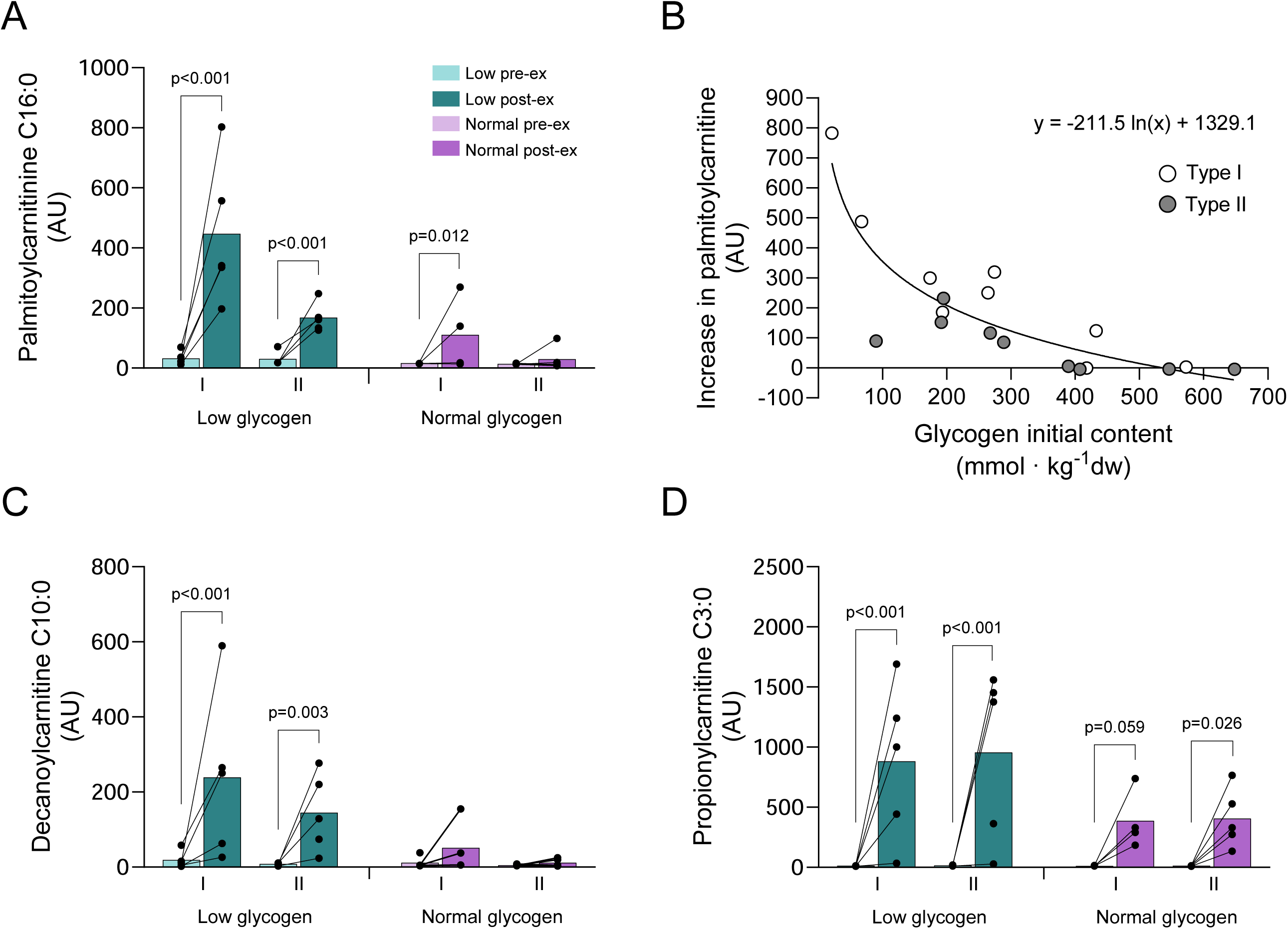
Fatty acids conjugated to carnitine in type I and type II fibers in the low and normal glycogen leg pre and post 60 min of two-legged cycling exercise. (A) palmitoylcarnitine, (B) increase in palmitoylcarnitine vs. initial muscle glycogen content, (C) decanoylcarnitine and (D) propionylcarnitine. Individual values from 5 subjects, except for type II fibers in the low leg pre-exercise (n=4) and type I fibers in the normal leg post- exercise (n=4) as well as mean values are presented. Linear mixed effects models were employed to identify differences between legs, fiber type and time.

The levels of glucose-6-phosphate and fructose-6-phosphate did not change significantly in any leg or fiber type during exercise, although in the normal leg, both metabolites increased in 4 of the 5 subjects in the type II fibers (Supplementary figure 2A, B).

The level of acetylcarnitine increased in both fiber types in the normal leg, 7 to 10-fold during exercise (P<0.001 for both fiber types), with a corresponding decrease in carnitine (P<0.001). In the low leg, there was an increase in acetylcarnitine (P<0.01) as well as a decrease in carnitine only in the type I fibers (P<0.05). The level of acetylcarnitine post exercise was higher in type I than in type II fibers in both legs (P<0.01), whereas that of carnitine was lower in type I fibers post exercise (P<0.05) only in the low leg (Supplementary figure 2C, D).

#### Cell signaling

The cycling exercise did not affect mTOR^Ser2448^ phosphorylation in the low leg but led to an increase in the normal leg, 2.9- and 2.3-fold increase in type I and type II fibers, respectively (P<0.05, Figure 7A). The increase in mTOR^Ser2448^ phosphorylation correlated significantly with the initial muscle glycogen content in type II fibers (r=0.81, P<0.05), but not in type I fibers (r=0.33, n.s.). The phosphorylation of 4E-BP1^Thr37/46^ decreased by 60-70% in both fiber types in both legs (P<0.05 for time) (Figure 7B), whereas the total protein remained unchanged in the normal leg but decreased by 33% (P<0.01) in both fiber types in the low leg (Supplementary figure 3C). The phosphorylation of eEF2^Thr56^ increased by 145% in type I fibers in the low leg (P<0.01), but there was no effect on this fiber type in the normal leg or in the type II fibers of either leg (Figure 7C). The change in eEF2^Thr56^ phosphorylation in type I fibers was negatively correlated to the initial content of muscle glycogen (Figure 7D), while no such correlation was found for type II fibers.

**Figure 7.**
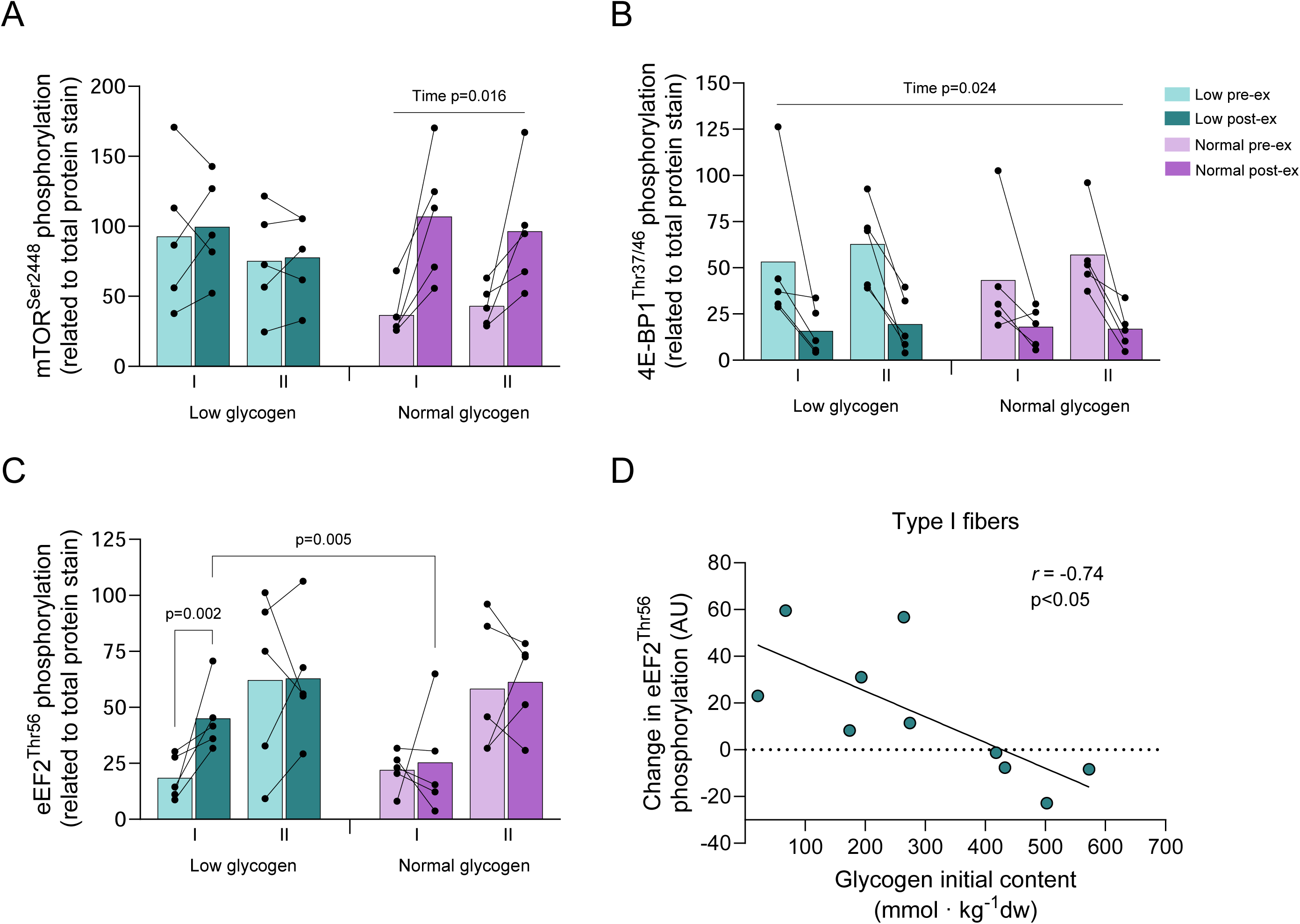
Protein phosphorylation (p) of (A) mTOR, (B) 4EBP-1, and (C) eEF2in type I and type II fibers in the low and normal glycogen leg pre and post 60 min of two-legged cycling exercise. Individual values from 5 subjects as well as mean values are presented. The ANOVA revealed a significant interaction between leg and time (p-mTOR) as well as an interaction between leg, fiber type and time (p-eEF2). Fisheŕs LSD post hoc test was employed to identify where the differences occurred. Changes in eEF2 phosphorylation vs. initial muscle glycogen content in type I fibers are shown in D.

The phosphorylation of AMPK^Thr172^ tended to increase, and the phosphorylation of ULK1^Ser555^ was significantly elevated post-exercise in both fiber types in both legs (P<0.05 for time), with no significant difference between fiber types or legs (Figure 8A, B). The phosphorylation of ACC^Ser79^ increased significantly during exercise in both legs and both fiber types (P<0.05) (Figure 8C).

**Figure 8.**
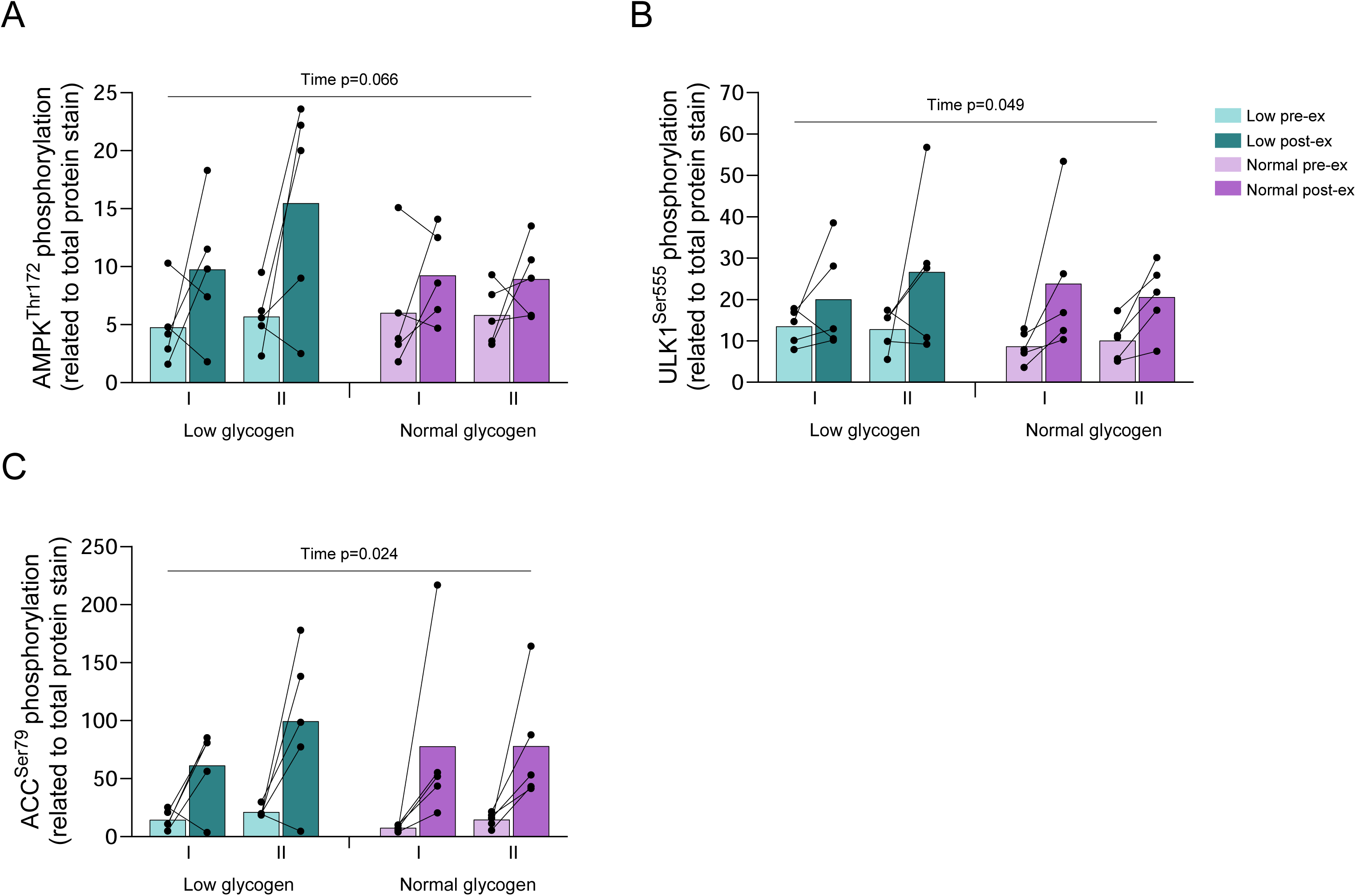
Protein phosphorylation (p) of (A) AMPK, (B) ULK-1, and (C) ACC in type I and type II fibers in the low and normal glycogen leg pre and post 60 min of two-legged cycling exercise. The ANOVA revealed a significant main effect of time for p-ACC and p-ULK-1.

The level of LC3B-I tended to decrease in the low leg (P=0.056 for time), mainly driven by a decrease in the type II fibers of all five subjects, but there was no change in any fiber type in the normal leg during exercise (Figure 9A). The decrease in the level of LC3B-II was numerically larger in both fiber types in the normal compared to the low leg (80% vs. 40% in type I and 90% vs. 70% in type II) and the level was lower in the normal compared to the low leg post-exercise (P<0.01 for pre- vs. post-exercise in both legs and P<0.05 for low vs. normal leg post-exercise, Figure 9B). Hence, the cycling exercise led to a significant reduction in the LC3B-II/LC3B-I ratio only in the normal leg (P<0.01). The ratio was reduced by 75% in type I fibers and 87% in type II fibers, with no significant difference between fiber types (Figure 9C). The decrease in LC3B-II content during exercise was inversely related to the increase in palmitoylcarnitine in such way that the largest decrease occurred when the increase of palmitoylcarnitine was small (Figure 9D). Exercise did not significantly influence the protein content of MuRF-1 in any fiber type or leg, although the content was significantly higher in both type I and type II fibers in the low leg (Supplementary figure 3I).

**Figure 9.**
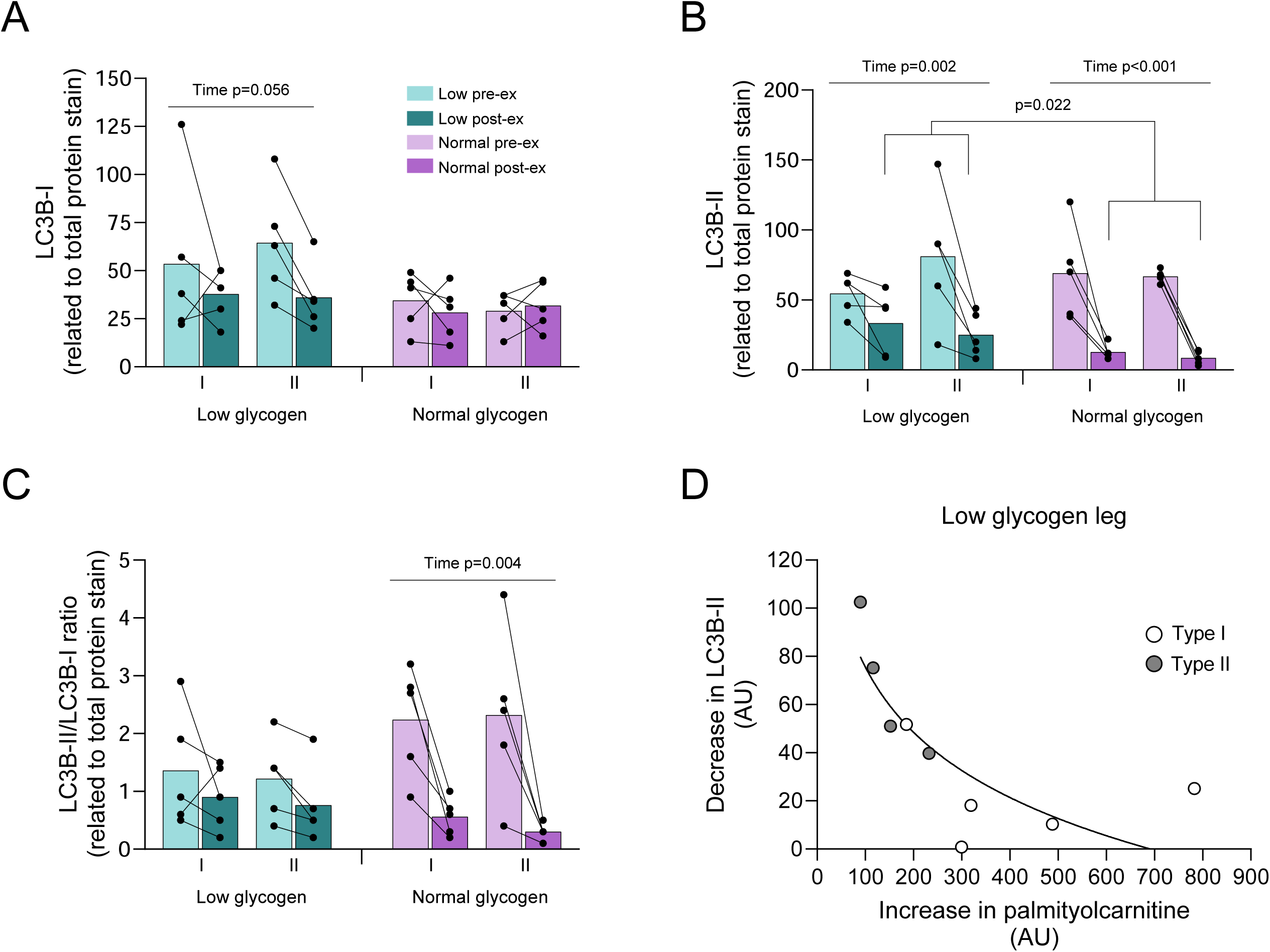
Protein content of (A) LC3B-I, (B) LC3B-II and (C) LC3B-II/I ratio in type I and type II fibers in the low and normal glycogen leg pre and post 60 min of two-legged cycling exercise, and (D) the relationship between the decrease in LC3B-II and the level of post-exercise palmitoylcarnitine in the low glycogen leg. Individual values from 5 subjects as well as mean values are presented. The ANOVA revealed a significant interaction between leg and time for all variables. Fisheŕs LSD post hoc test was employed to identify where the differences occurred.

The total protein content of mTOR, S6K1, eEF2, AMPK, ACC, ULK1, and UBR5 was similar in both fiber types and legs before exercise, and none of the proteins was significantly affected by exercise on day 2 (Supplementary figure 3).

## Discussion

We investigated the effect of exercise with low glycogen availability on metabolism and cell signaling in a large number of type I and type II fibers. The study design enabled us to evaluate the effect of glycogen content independent of hormonal changes and the delivery of blood-borne substrates. The primary and novel findings are that 1) the utilization rate of muscle glycogen during exercise was reduced for both fiber types in the low compared to the normal leg, 2) carnitine-conjugated long-chain fatty acids were 4-5 times higher in both type I and type II fibers in the low compared to the normal leg post-exercise, 3) phosphorylation of mTOR^Ser2448^ increased in the normal leg but remained unchanged in both fiber types in the low leg together with increased phosphorylation of eEF2^Thr56^ only in type I fibers post-exercise, and 4) the level of LC3B-I tended to decrease during exercise only in the low leg, whereas LC3B-II decreased in both legs but to a lower level in the normal leg post exercise, causing a reduction in the LC3B-II/I ratio in both fiber types only in the normal leg.

The observed reduced rate of glycogen utilization in both fiber types in the low leg is in agreement with studies using mixed muscle, where the glycogen utilization rate is associated with the initial content of muscle glycogen [36, 37]. In addition, the present study shows that while this holds for both fiber types, it is most obvious for the type I fibers where a highly significant correlation was found between initial muscle glycogen levels and the utilization of glycogen during exercise (Figure 5B).

Carnitine is essential for the transport of long-chain fatty acids from the cytosol into the mitochondria for subsequent oxidation, and recent data indicate that carnitine also plays a role in the utilization of medium-chain fatty acids in skeletal muscle [38]. However, while we did not measure lipid oxidation *per se*, our findings that carnitine-conjugated fatty acids were elevated to much larger extent in the low leg post-exercise, likely reflect a greater breakdown and utilization of fatty acids in both fiber types in this leg. A shift towards increased lipid oxidation when the glycogen level is low is mediated by, among others, changes in hormone and substrate levels that favor uptake of fatty acids by muscle [39]. In the present study, both legs were exposed to the same arterial concentration of hormones and fatty acids. Our results therefore indicate a greater utilization of intramuscular triglycerides rather than blood-borne fatty acids. This is further supported by the observation that only minor or no differences in uptake of fatty acids were detected between the normal and a low glycogen leg in previous studies with a similar design [5, 30].

It is well-documented that type I fibers have a higher content of triglycerides as well as a higher capacity to oxidize fat [40–44], and therefore utilize more intramuscular triglycerides than type II fibers during exercise [40, 44, 45]. In line with this, the increase in carnitine-bound long-chain fatty acids was more than twice as high in the type I compared to the type II fibers in the low leg, and furthermore the level was increased only in type I fibers in the normal leg (Figure 6). The larger increase in both fiber types in the low leg may be expected considering that the rate of glycogen utilization in the type I and type II fibers in this leg was only 20% and 45%, respectively, of the rate in the normal leg, although this has not been reported previously.

When exercise began with low levels of muscle glycogen the phosphorylation of mTOR^Ser2448^ was unaffected and 4E-BP1^Thr37/46^ phosphorylation decreased in both fiber types while eEF2^Thr56^ phosphorylation increased only in the type I fibers. Phosphorylation of eEF2 at Thr56 is known to inactivate the enzyme [46], and possibly also translation elongation and the subsequent rate of protein synthesis. The inverse correlation between the increase in eEF2^Thr56^ phosphorylation and the level of muscle glycogen in type I fibers suggests a regulatory role of glycogen (Figure 7D). Moreover, it appears that phosphorylation of eEF2^Thr56^ is induced only when the glycogen level falls below 300 mmol kg^-1^ dw. Interestingly, this is also the level of glycogen below which the palmitoylcarnitine begins to increase during exercise (Figure 6B). Accordingly, either a direct or an indirect effect of low muscle glycogen may cause an inhibition of mTORC1 (see below). Overall, our data suggest that during exercise with limited availability of glycogen, the rate of protein synthesis is suppressed in both fiber types but possibly even more so in type I fibers. Some support for a selective inhibition of the protein synthetic rate in type I fibers is presented in a study by Rose and coworkers (2009). Using immunohistochemical staining of muscle cross-sections, they observed an increased eEF2 phosphorylation in type I fibers following high-intensity exercise [11]. Furthermore, resistance exercise with low levels of glycogen has been reported to inhibit mTORC1 signaling in mixed muscle [47], and although the design and type of exercise were different from the present study, our results are in line with this observation.

The marked increase in the levels of fatty acids conjugated to carnitine in the low leg post-exercise coincided with a blunted mTORC1 signaling response, more apparent in type I than type II fibers through increased eEF2^Thr56^ phosphorylation. It is therefore possible that fatty acids have a direct inhibitory effect on the mTORC1 signaling cascade. Mechanistically, this is supported by work *in vitro* showing that palmitic acid impairs S6K1^Thr389^ phosphorylation in C2C12 cells [48], and several studies have reported attenuated muscle adaptation to exercise during a high-fat diet [49]. The impaired mTORC1 activation in the present study may therefore represent an inhibitory effect of increased fatty acids as a consequence of a metabolic switch during conditions of low glycogen availability.

The phosphorylation of eEF2^Thr56^ was considerably higher in type II than in type I fibers in resting muscle in agreement with previous studies [11, 50], suggesting a higher basal rate of protein synthesis in type I than in type II fibers [51]. A higher rate of protein synthesis in slow-twitch compared to fast-twitch muscle has previously also been reported in rodents [52, 53]. Whether a possible fiber-type specific difference in resting synthetic rate may influence the response to exercise with reduced availability of muscle glycogen is presently unknown.

Endurance exercise is considered to stimulate autophagy, and several studies have indicated that induction of autophagy is essential for muscular adaptation to exercise training [17]. During autophagy, lipidation of LC3-I forms LC3B-II located at the autophagosome membrane. The level of LC3B-II is therefore frequently used as a marker for autophagosome content and also an indirect marker of autophagic activity (see Botella et al. 2024) [54]. Endurance exercise may regulate autophagy in different ways. One is through activation of mTORC1, which is a regulator not only of the rate of protein synthesis but is also a key regulator of the autophagy pathway in skeletal muscle. Activation of mTORC1 negatively regulates autophagic induction by phosphorylating ULK1 on the Serine 757 site [55].

In the normal leg, exercise caused an increase in mTOR^Ser2448^ phosphorylation, a pronounced reduction in LC3B-II content, as well as a decrease in the ratio LC3B-II/LC3B-I in both fiber types. These findings are in agreement with previous studies on mixed muscle, suggesting reduced autophagosome formation [17, 54, 56, 57]. On the contrary, for the low leg, exercise did not affect mTOR^Ser2448^ phosphorylation, the reduction in LC3B-II was smaller than seen in the normal leg, and the LC3B-II/LC3B-I ratio remained essentially unchanged thus suggesting an attenuated effect. Due to limited sample availability and antibody issues, we were unable to quantify ULK1^Ser757^. Interestingly, we found a greater reduction in LC3B-II content when the increase in palmitoylcarnitine level was small which coincides with the availability of muscle glycogen being sufficient (Figures 6B and 9D).

The energy sensor AMPK is known to regulate autophagy through direct activation of ULK-1 on the Serine 555 site. However, the expected differential activation of these proteins in the low and normal leg was not detected. Instead, exercise caused similar increases in phosphorylation of AMPK^Thr172^ and ULK-1^Ser555^ in both legs as well as in both fiber types (Figure 9). Our data thus indicate that mTORC1 signaling is more important in regulating the autophagic response in this specific exercise situation, as was suggested to be the case for the autophagy-inhibiting effect of insulin [57]. The role of mTORC1 in regulating autophagy was recently confirmed in a study on different fiber types following ingestion of a mixed meal [58]. Here, the observed reduction in LC3B-II abundance in both type I and type II fibers occurred together with similar increases in Akt and mTOR phosphorylation [58].

We found no fiber type-specific difference in LC3B-II content, in contrast to studies on rodents where autophagy protein content and flux were higher in slow-twitch than in fast-twitch muscle [59]. In agreement with this, a higher level in type I versus type II fibers was recently reported in human subjects [58]. The reason for the divergent results is unclear, but may be related to the subject groups. In the study by Morales-Scholz and coworkers (2022) the subjects were sedentary overweight in contrast to our group of moderately trained subjects, who may have a relatively high oxidative capacity also in their type II fibers [60].

Twelve hours after the glycogen reduction exercise on day 1, the level of mTORC1^Ser2448^ phosphorylation was higher in both fiber types, suggesting that activation of this protein persists despite no nutritional intake. The upregulation of MuRF-1 expression in both fiber types, while larger in type II fibers, and the elevation of the autophagy protein LC3B-I in both fiber types, suggest an adaptation to training and possibly increased capacity in the degradation pathways. However, to what extent the effects of the one-legged exercise on day 1 influenced the response to exercise on day 2 is not known. Based on the inverse relationships between glycogen level and the phosphorylation of eEF2^Thr56^ in type I fibers and carnitine-fatty acids and reduction in the LC3B-II content in the low leg, it appears more likely that the attenuated response is related to the low initial glycogen levels.

## Conclusions

Our findings show that when exercise begins with low levels of muscle glycogen, phosphorylation of mTOR^Ser2448^ remains unchanged in both type I and type II fibers, and eEF2^Thr56^ increases only in type I fibers indicating that the rate of protein synthesis is depressed primarily in these fibers. In addition, exercise induced only minor or no reduction in the level of autophagic markers in both fiber types in the leg with low glycogen level, possibly due to the absence of mTOR activation. This suggests that exercise with limited availability of glycogen may prevent the typically observed reduction in autophagy. Furthermore, these effects appeared to be driven by changes in substrate availability and/or utilization during exercise rather than the remaining effect of the glycogen reduction exercise performed a day earlier.

## Declarations

### Ethics approval and consent to participate

The study was approved by the Swedish Review Authority (2018/2186-31) and performed in accordance with the principles outlined in the Declaration of Helsinki. All participants were fully informed about the experimental procedure and associated risks before giving their written consent.

## Consent for publication

Not applicable

## Availability of data and materials

All data generated or analyzed during the study are available from the corresponding author upon request.

## Authorś contributions

EB and SE conceived and designed research; EB, SE and OH performed experiments; LC, HS, MM, KS, AC and EB performed analyses; EB, OH, MM, JLR and KS analyzed data and interpreted results; OH prepared figures; EB, OH drafted manuscript; MM, JLR, SE, LC and KS edited and revised manuscript. All authors read and approved the final manuscript.

## Acknowledgments

The authors wish to thank the participants for their time and effort. The authors also wish to thank Professor Björn Ekblom and Professor Lars Larsson for their assistance during sample collection. We acknowledge the Karolinska Institute Small Molecule Mass Spectrometry Core Facility (KI-SMMS), supported by KI/SLL, for support in the sample analyses.

---

**Supplementary figure 1.**
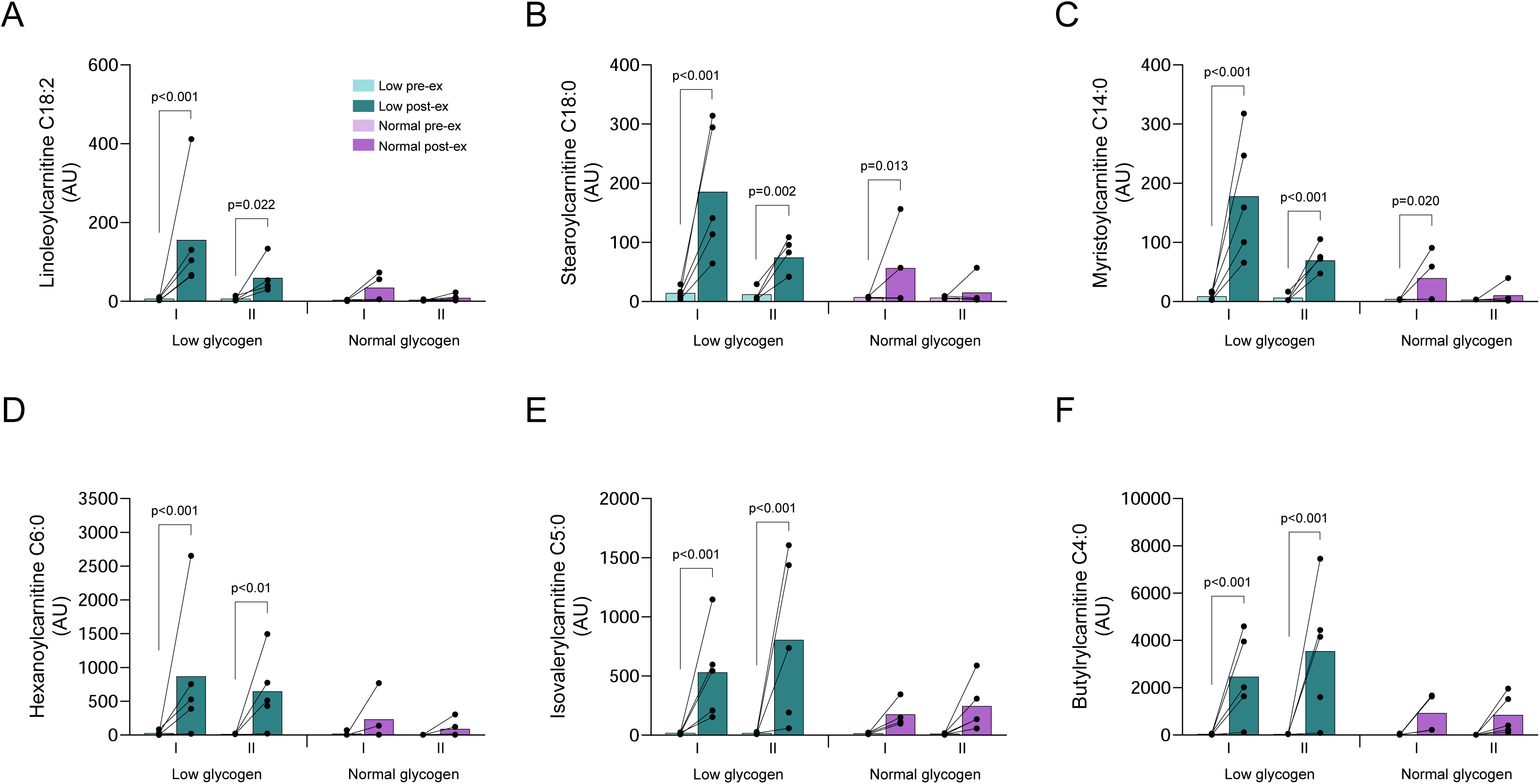
Levels of fatty acids conjugated to carnitine in type I and type II fibers in the low and normal glycogen leg pre and post 60 min of two-legged cycling exercise. (A) linoleoylcarnitine, (B) stearoylcarnitine, (C) myristoylcarnitine, (D) hexanoylcarnitine, (E) isovalerylcarnitine and (F) butyrylcarnitine. Individual values from 5 subjects, except for type II fibers in the low leg pre-exercise (n=4) and type I fibers in the normal leg post-exercise (n=4), as well as mean values are presented. Linear mixed effects models were employed to identify differences between legs, fiber type and time.

**Supplementary figure 2.**
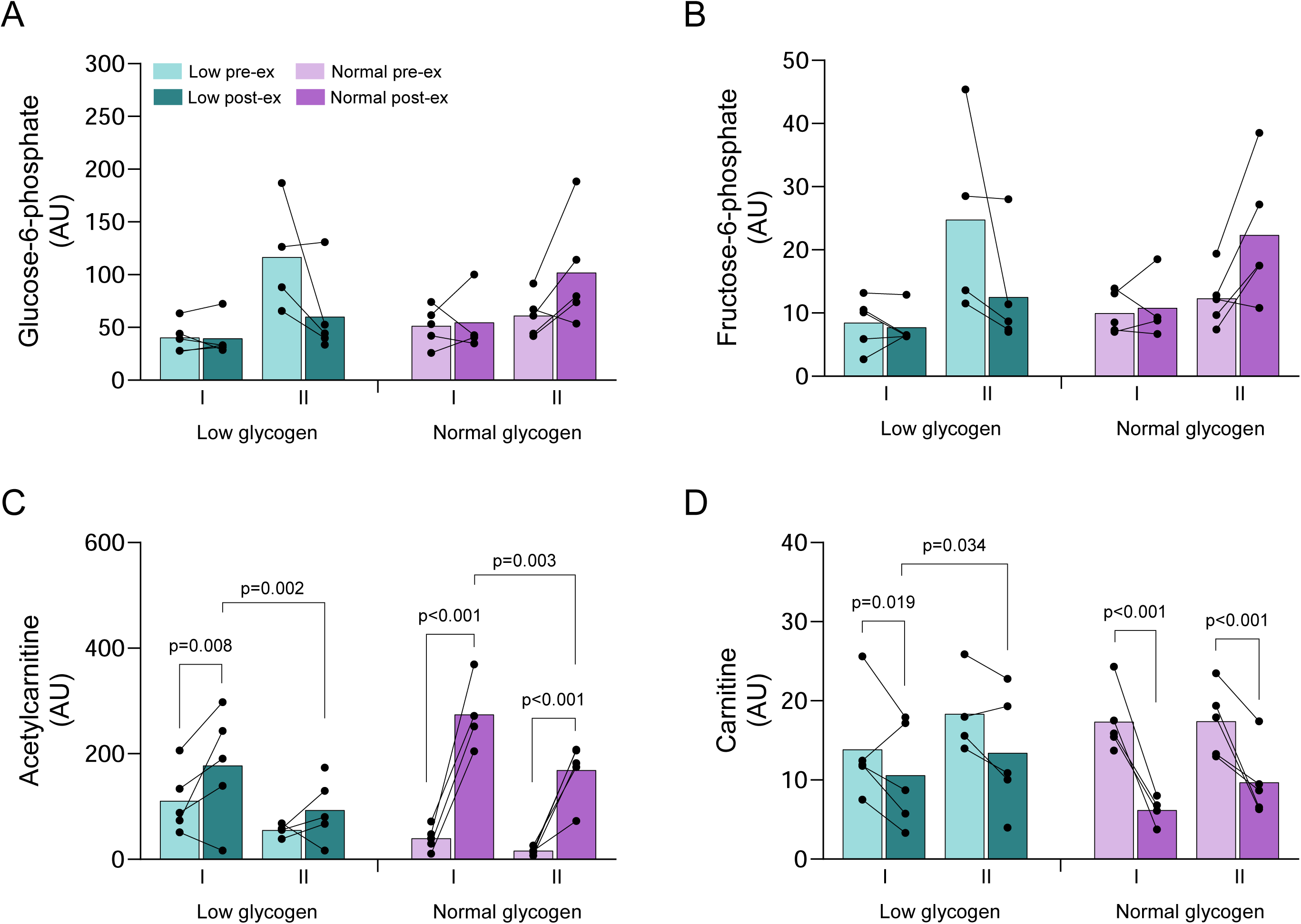
(A) glucose-6-phosphate, (B) fructose-6-phosphate, (C) acetylcarnitine and (D) carnitine in type I and type II fibers in the low and normal glycogen leg pre and post 60 min of two-legged cycling exercise. Individual values from 5 subjects as well as mean values are presented except for type II fibers in the low leg pre-exercise (n=4) and type I fibers in the normal leg post-exercise (n=4). Linear mixed effects models were employed to identify differences between legs, fiber type and time.

**Supplementary figure 3.**
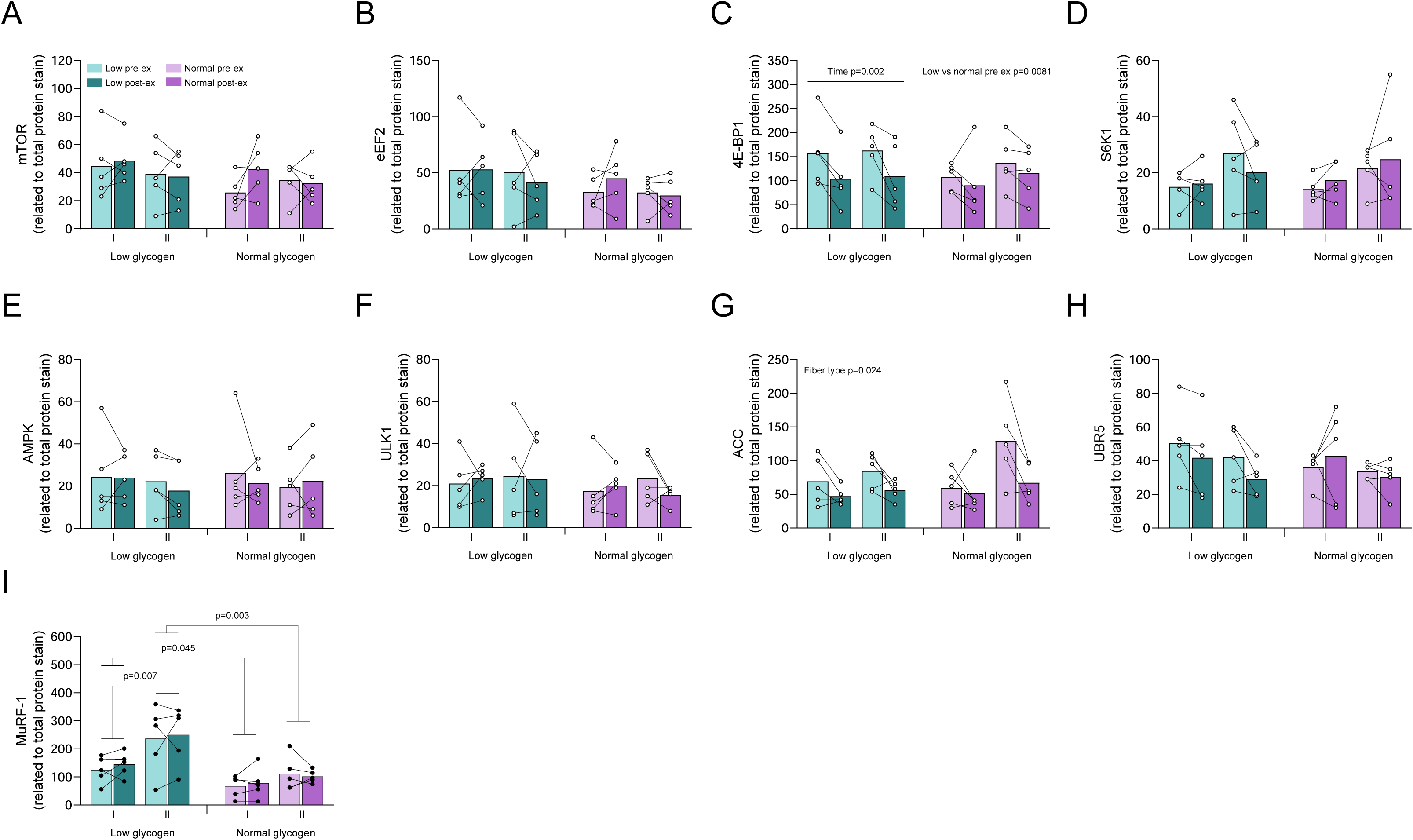
Protein content of (A) mTOR, (B) eEF2, (C) 4EBP-1, (D) S6K1, (E) AMPK, (F) ULK1, (G) ACC and (H) UBR5 in type I and type II fibers in the low and normal glycogen leg before and after 60 min of cycling exercise. Individual values from 5 subjects as well as mean values are presented. The ANOVA revealed a significant interaction between leg and time for 4E-BP1 content, and the Fisheŕs LSD post hoc test revealed a difference between the low and normal leg pre-exercise and between pre and post-exercise for the low leg.

**Supplementary figure 4.**
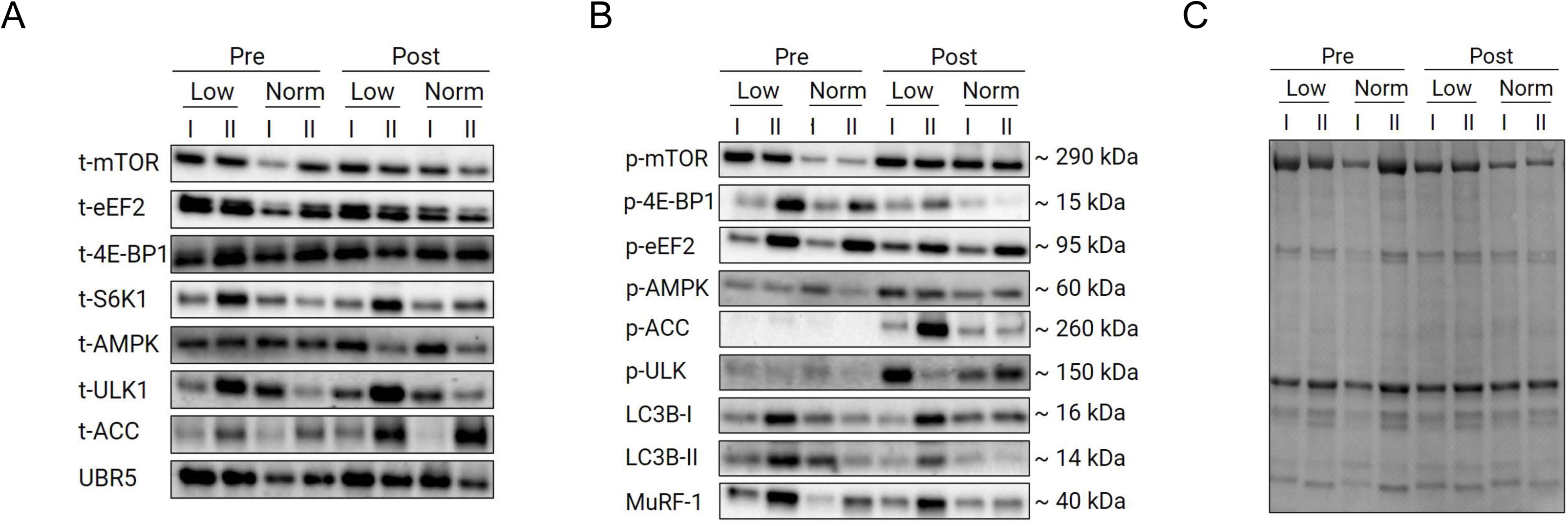
Representative immunoblots of total protein content (A), phosphorylated protein (B) and total protein stain (MemCode) (C).

